# Integrative analysis of nanopore direct RNA sequencing data reveals a role of PUS7-dependent pseudouridylation in regulation of m^6^A and m^5^C modifications

**DOI:** 10.1101/2024.01.31.578250

**Authors:** Mohit Bansal, Anirban Kundu, Anamika Gupta, Jane Ding, Andrew Gibson, Sanjay Varma RudraRaju, Sunil Sudarshan, Han-Fei Ding

## Abstract

Understanding the interactions between different RNA modifications is essential for unraveling their biological functions. Here, we report NanoPsiPy, a computational pipeline that employs nanopore direct RNA sequencing to identify pseudouridine (Ψ) sites and quantify their levels at single-nucleotide resolution. We validated NanoPsiPy by transcriptome-wide profiling of PUS7-dependent Ψ sites in poly-A RNA and rRNA. NanoPsiPy leverages Ψ-induced U-to-C basecalling errors in nanopore sequencing data, allowing detection of both low and high stoichiometric Ψ sites. We identified 8,624 PUS7-dependent Ψ sites in 1,246 mRNAs encoding proteins associated with ribosome biogenesis, translation, and energy metabolism. Importantly, integrative analysis revealed that PUS7 knockdown increases global mRNA N^6^-methyladenosine (m^6^A) and 5-methylcytosine (m^5^C) levels, suggesting an antagonistic relationship between Ψ and these modifications. Our study underscores the potential of nanopore direct RNA sequencing in revealing the co-regulation of RNA modifications and the capacity of NanoPsiPy in analyzing pseudouridylation and its impact on other RNA modifications.

## Introduction

Research spanning several decades has demonstrated that RNAs undergo various posttranscriptional modifications, including pseudouridylation^1–3^. Pseudouridine (Ψ) is the first identified RNA modification^4^ that constitutes 0.2–0.6% of total uridines in mammalian mRNAs^5^. Ψ can enhance the stability and rigidity of RNAs, influencing cellular processes such as translation, splicing, and telomere maintenance^2,6^. The existence of Ψ in mRNAs remained unknown until 2014-2015^5,7,8^, a revelation facilitated by advancements in next-generation sequencing technologies. Human genome encodes thirteen annotated pseudouridylation installation enzymes (pseudouridine synthases, PUSs)^1^, highlighting the robust and dynamic nature of Ψ installation in human RNA. The conversion of uridine to Ψ starts with breaking the N1-C1’ bond linking the uracil base to the sugar, followed by an 180° base rotation around the N3-C6 axis. The resulting Ψ contains an extra hydrogen bond donor at N1 and a C5-C1’ base- sugar linkage^9^. The success of mRNA-based COVID-19 vaccines in which all uridines are replaced by Ψs underscores the crucial role played by Ψ modification in promoting protein expression and increasing the efficacy of mRNA vaccines, showcasing the importance of understanding and harnessing RNA modifications for therapeutic purposes^10^. Ψ does not disrupt canonical base pairing, making it indistinguishable from uridine in hybridization-based methods. Additionally, the molecular weight of Ψ matches that of uridine, complicating direct detection through mass spectrometry^11^. This difficulty is exacerbated when trying to identify Ψ sites within specific mRNA sequences.

Various techniques, such as Pseudo-Seq^7^, Ψ -Seq^8^, and Ceu-Seq^5^ that are based on illumina-short read sequencing, have used the CMC (N-cyclohexyl-N’-β-(4-methylmorpholinium)ethylcarbodiimide)-based approach for transcriptome-wide Ψ mapping. While CMC-based methods are effective in generating a stop signature during reverse transcription (RT) for Ψ site identification, these methods lack base resolution and stoichiometry information at the modified bases. In contrast to aforementioned techniques, a new generation of methods based on bisulfite-induced deletion signatures at Ψ sites, including RBS-seq^12^, BID-seq^13^, and PRAISE^14^, offer quantitative transcriptome-wide mapping of Ψ sites. However, it is important to note that Ψ-bisulfite adduct formation involves harsh chemical reactions as an intermediate step. Moreover, the read-length limitation of Illumina sequencing also narrows the possibility to examine Ψ distribution in mRNA splice isoforms and the linkage of multiple Ψ sites in single RNA molecules.

We wanted to develop a nanopore direct RNA sequencing method for Ψ site identification and stoichiometry measurement on long native mRNAs without reliance on harsh chemical reactions, providing a more direct and precise means of studying pseudouridylation. Several studies have utilized nanopore-based direct RNA sequencing to directly examine RNA modifications. The detection of Ψ using direct RNA sequencing has been confirmed for ribosomal RNAs (rRNAs) in Saccharomyces cerevisiae^15^ and human mRNA^16,11^, based on U-to-C basecalling errors. However, it remains to be established whether direct RNA long-read nanopore sequencing is sufficiently robust to determine PUS-dependent regulation of Ψ location and stoichiometry in mRNA. This study aimed to develop a method to validate the U-to-C basecalling signature as a distinctive feature of Ψ and to quantify transcriptome-wide PUS7-dependent Ψ sites using in vivo and in vitro systems. Furthermore, cross talk between RNA modifications can significantly impact mRNA expression and translation, highlighting the intricate regulatory networks within cells. Consequently, we investigated the interplay between Ψ, N^6^-methyladenosine (m^6^A), and 5-methylcytosine (m5C) to better understand their interactions.

## Results

### Development of NanoPsiPy for identification and quantification of Ψ modification based on U-to-C basecalling errors in nanopore sequencing

We have developed NanoPsiPy, a computational pipeline (https://github.com/vetmohit89/NanoPsiPy), to identify and quantify Ψ levels in RNA based on U-to-C basecalling errors in Nanopore direct RNA sequencing data (Figure 1A). Following the alignment of RNA reads to the human reference sequence, NanoPsiPy systematically gathers positional data for each uridine base. Subsequently, it conducts statistical comparisons of U-to-C basecalling errors between two sample groups (control and treatment), identifying and quantifying Ψ modifications at specific locations. As Ψ detection by Nanopore direct RNA sequencing is based on basecalling errors, it is essential to develop a method that accounts only primary alignment of each RNA read (i.e., the alignment with the highest mapping quality) to reduce false positive errors. NanoPsiPy filters out secondary alignments and only uses reads with mapping quality score of 20 or higher, which correspond to the probability of correct alignment of 99% or higher, for downstream analysis and calculating U-to-C basecalling errors.

**Figure 1.**
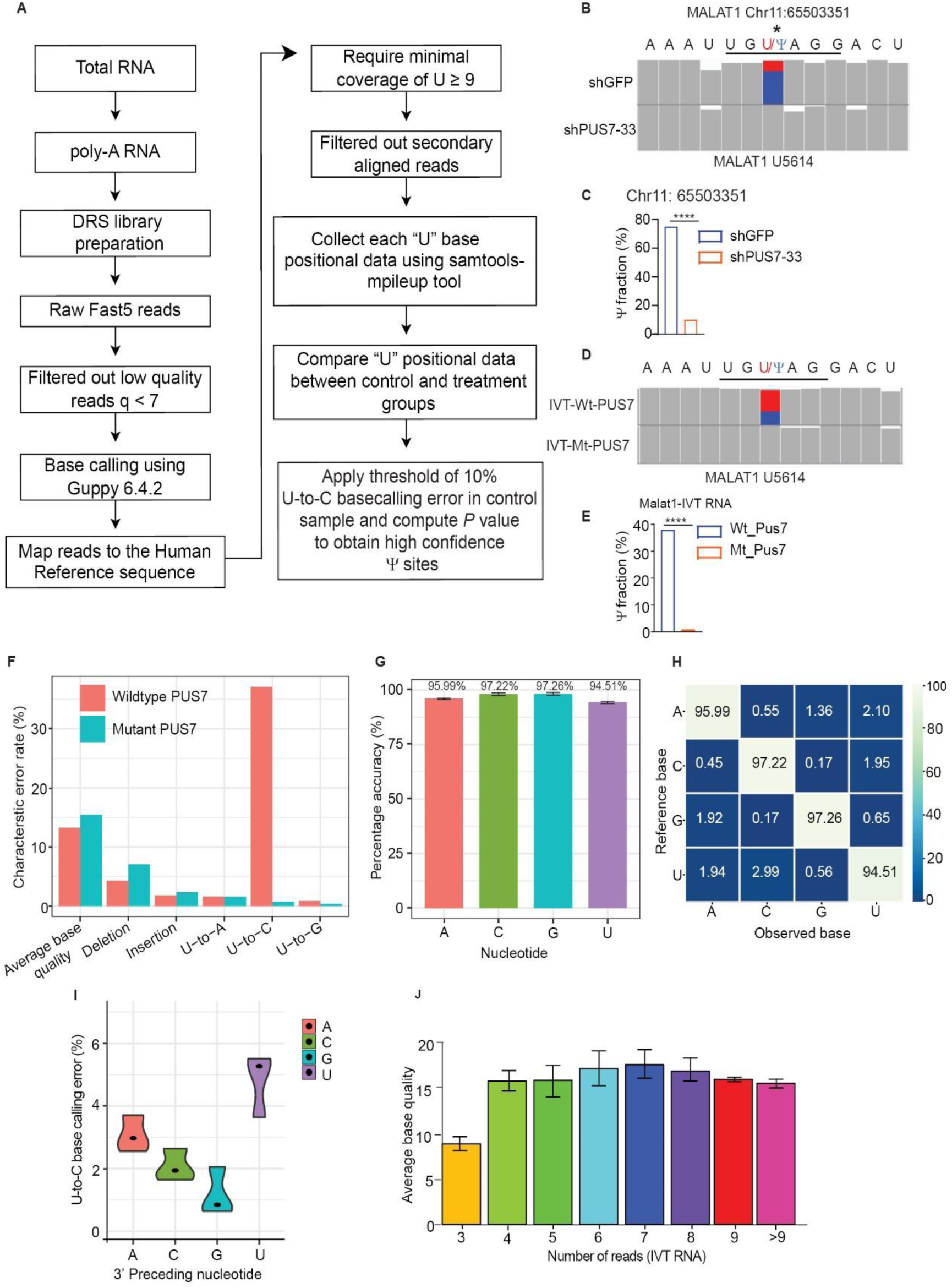
Identification and quantification of **Ψ** based on NanoPsiPy analysis of nanopore sequencing data and U-to-C basecalling errors. **(A)** Flow chart for using the NanoPsiPy pipeline to detect and quantify Ψ based on nanopore direct RNA sequencing. **(B)** Integrated Genome Viewer (IGV) snapshots of aligned nanopore MALAT1 RNA from BE(2)-C shGFP and BE(2)-C shPUS7-33 cells. **(C)** U-to-C mismatch percentage for U5614 based on data from **(B**). U5614 is shown as red and Ψ as blue, and correctly aligned bases are shown in gray. Genomic reference sequence is converted to the sense strand and shown as RNA for clarity. **(D)** IGV snapshots of aligned nanopore MALAT1 IVT RNA oligo treated with immunopurified wildtype or mutant PUS7. **(E)** U-to-C mismatch percentage for U5614 based on data from **(D**). **(F)** Comparison of basecalling features between the samples treated with immunopurified wild-type or mutant PUS7. **(G-H)** Accuracy of base-calling using a pool of three different IVT RNA oligos. **(I)** Effect of different preceding bases on estimating U- to-C basecalling errors using a pool of three different IVT RNA oligos. **(J)** Average base quality scores with respect to the number of reads for IVT RNA samples. ****P < 0.0001.

Another potential source of error is the presence of single-nucleotide polymorphisms (SNPs) whereby the base is different from the reference genome or transcriptome sequence. SNPs can cause systematic basecalling errors. NanoPsiPy removes any SNP errors from downstream analysis by comparative analysis of the U-to-C mismatch at each site between control and treatment samples (Figure 1A).

We tested the ability of our nanopore direct RNA sequencing and the NanoPsiPy pipeline to identify and quantify Ψ based on U-to-C basecalling errors using two PUS7- dependent systems. We used lentiviruses expressing a shRNA against human PUS7 (shPUS7-33) to silence PUS7 expression in the *MYCN*-amplified neuroblastoma cell line BE(2)-C, with shGFP as control (Supplemental Figure 1). It has been shown previously that PUS7 pseudouridylates U5614 in MALAT1 RNA (base position according to NR-002819.4)^17^. In control shGFP BE(2)-C cells, MALAT1 RNA at U5614 showed 75% of RNA reads having a U-to-C basecalling error, which was reduced to 10% in BE(2)-C cells with PUS7 knockdown (Figures 1B-C). To further validate this U- to-C basecalling error as a signature of PUS7-dependent Ψ modification, we performed an in vitro pseudouridylation assay^18^ using an in vitro transcribed (IVT) MALAT1 RNA oligo that includes 86 nucleotides upstream and 66 nucleotides downstream of U5614. The RNA oligo was incubated either with immunopurified wild-type human PUS7 or with a catalytically inactive mutant (PUS7 D294A)^19^ and analyzed systematically for basecalling features, including mismatches, deletion, insertion and per-base qualities.

Approximately 38% of the IVT MALAT1 RNA reads treated with wild-type PUS7 showed a U-to-C basecalling error compared to the 1% error in the RNA reads treated with the mutant PUS7 (Figures 1D-E). We found no significant differences in other basecalling features between the two samples (Figure 1F). These findings confirmed that NanoPsiPy analysis of nanopore direct RNA sequencing data was able to detect and quantify Ψ in RNA by identifying U-to-C basecalling errors.

Nanopore sequencing detects nucleotide bases from their ionic current signature.

Our pipeline NanoPsiPy uses Guppy (v6.4.2) for basecalling. By using a pool of different IVT RNA oligos containing only canonical RNA nucleotides, we calculated the basecalling accuracy. We found that our pipeline is highly accurate, with 95.99% for adenine, 97.22% for cytosine, 97.26% for guanine, and 94.51% for uridine (Figure 1G). Additionally, we found that Guppy could reliably identify unmodified and aligned uridine with an average background U-to-C calling error of 2.99% (Figure 1H).

Nanopore direct RNA sequencing method sequences RNA in the 3’ to 5’ direction, raising the question of whether U-to-C error-based Ψ detection is impacted by the preceding 3’ base. Our analysis of IVT RNA data revealed that when the preceding 3’ base is U, A, C, or G, there is a background error of 4.8%, 3.07%, 2.07%, or 1.18% in identifying the candidate U as C, respectively (Figure 1I). Further, we observed that the average base quality is low when RNA reads at any location are less than 4 (Figure J). Therefore, to minimize the inclusion of U sites with background errors, we defined a threshold of 10% basecalling error for U-to-C estimation and a minimum of 9 reads at any given U site for Ψ detection and quantification in the whole transcriptome.

### Transcriptome-wide identification and quantification of Ψ sites in poly-A RNA from BE(2)-C cells using nanopore direct RNA sequencing and NanoPsiPy

We applied nanopore direct RNA sequencing and NanoPsiPy for de novo detection and quantification of transcriptome-wide Ψ modifications in the *MYCN*-amplified neuroblastoma cell line BE(2)-C. For background control (RNA without modifications), we used the published nanopore direct RNA sequencing data from a HeLa IVT poly-A RNA library that was constructed first by reverse transcription of HeLa poly-A RNAs (with modifications) into cDNAs to remove all modifications and then by in vitro transcription of the cDNAs back into RNAs^11^. We conducted nanopore direct sequencing of poly-A RNA from BE(2)-C shGFP cells and compared the sequencing data with the HeLa IVT poly-A RNA nanopore sequencing data, which showed an overlapping distribution of read lengths (Figure 2A). We then mapped RNA sequencing reads from both datasets to the human genome reference sequence (GRCh38.p14) to identify all the uridine sites common between BE(2)-C and HeLa IVT RNA and assigned the U-to-C basecalling error as Ψ modification at each site. We defined Ψ sites in BE(2)-C poly-A RNA based on two criteria: 1) the U-to-C basecalling error at each site in BE(2)-C must exceed 10% and 2) the *p* value of U-to-C basecalling error must be < 0.01. Using data from more than 2 million uridines in BE(2)-C poly-A RNA and more than 3 million uridines in HeLa IVT RNA, we identified 39,756 Ψ sites in BE(2)-C poly-A RNA with a median Ψ level of 16.67% (Supplemental Table 1). To assess the accuracy of our NanoPsiPy pipeline in detecting Ψ sites, we compared the identified BE(2)-C Ψ sites with the previously annotated 1,691 Ψ sites in HeLa by nanopore direct RNA sequencing^11^. Our analysis revealed 517 Ψ sites shared between BE(2)-C and HeLa cells (Supplemental Table 2). Furthermore, we observed a strong correlation (R² = 0.618) in the Ψ fraction across these shared Ψ sites (Figure 2B).

**Figure 2.**
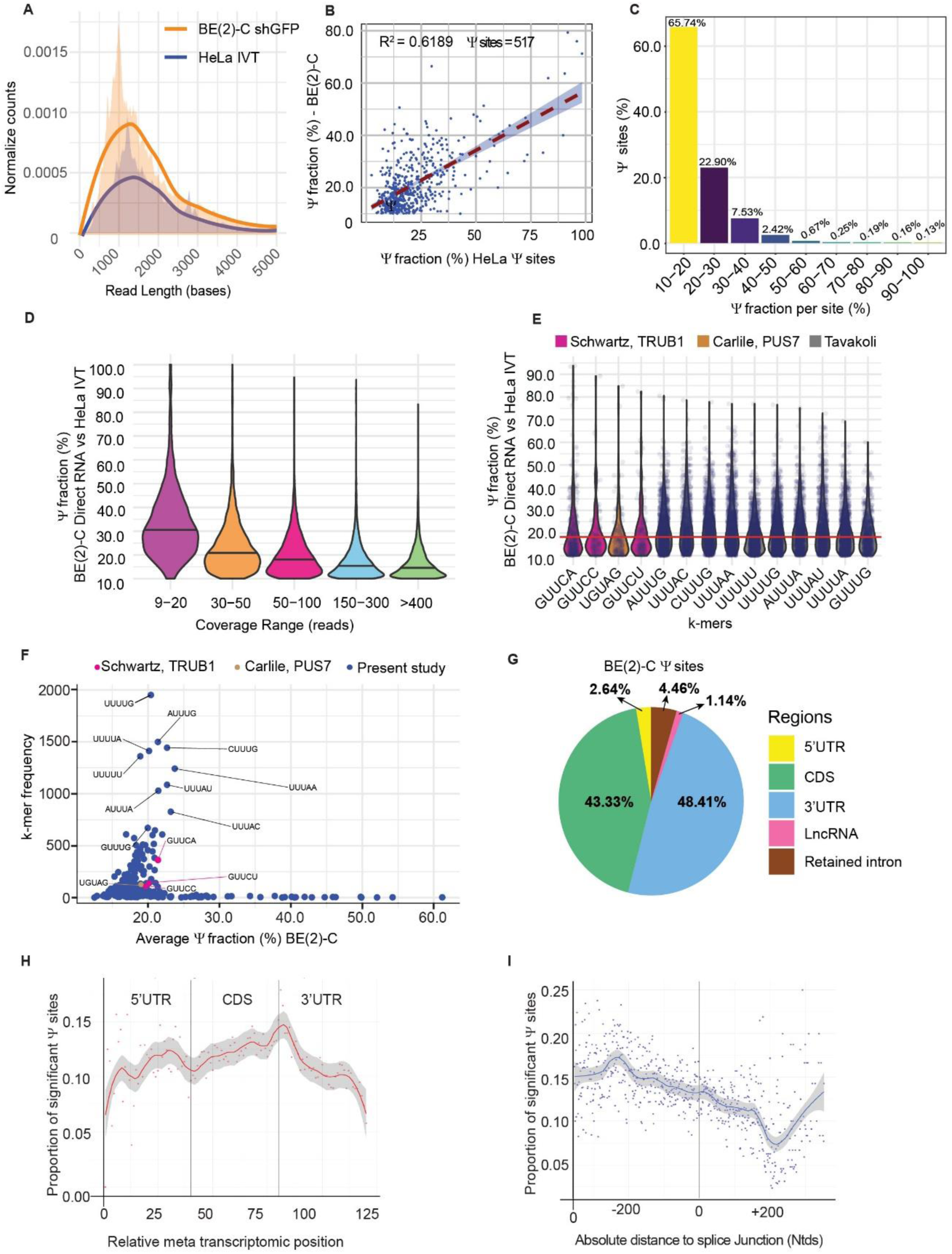
Transcriptome-wide identification and quantification of Ψ sites in poly-A RNA from BE(2)-C cells using nanopore direct RNA sequencing and NanoPsiPy. **(A)** Normalized counts of read lengths for BE(2)-C poly-A RNA (orange) versus HeLa IVT RNA (blue). (**B)** Scatter plot showing distribution of Ψ fractions (stoichiometries) of 517 Ψ sites shared by BE(2)-C and HeLa poly-A RNA. **(C)** Bar plot showing the proportional distribution of Ψ sites based on Ψ fractions (stoichiometries) for all Ψ sites identified in BE(2)-C poly-A RNA. **(D)** Bar plot showing the distribution of Ψ fractions in BE(2)-C poly-A RNA based on read coverages (reads). **(E)** K-mers frequency of most represented and frequently detected Ψ sites with high confidence. **(F)** K-mers frequency versus average Ψ fraction for all Ψ sites in BE(2)-C poly-A RNA. **(G)** Distribution of Ψ sites in mRNA regions (5′UTR, CDS and 3’UTR), long non-coding RNA, and retained intron. **(H)** MetaTranscript plot showing the proportion of Ψ along relative positions in the 5’UTR, CDS, and 3’UTR of BE(2)-C mRNA. (**I**) MetaJunction plot showing Ψ sites relative to their distances to the closest downstream (left) and upstream (right) exon- exon junction.

Next, we examined the stoichiometric distribution of Ψ levels in BE(2)-C poly-A RNA. Our analysis revealed that approximately 66% of the Ψ sites were modified at the levels of 10-20% and ∼11% at the levels of more than 30% (Figure 2C). In addition, our analysis revealed a negative correlation between the levels of Ψ modification and the levels of sequencing coverage (reads) at specific sites: the lower the number of reads, the higher the median Ψ levels (Figure 2D).

We next analyzed the sequence contexts for the BE(2)-C Ψ sites and found that the TRUB1 motifs GUUCA, GUUCC, and GUUCU, and the PUS7 motif UGUAG were the most highly represented (Figure 2E). The median level of Ψ across the top 14 motifs was 18.26% (Figure 2E). For all the Ψ sites, uridine exhibits a strong preference at the - 2, -1, and +1 positions (Figure 2F). The most frequent k-mers are UUUUG (4.92%), AUUUG (3.78%), UUUUA (3.56%), CUUUG (3.64%), and UUUUU (3.43%) (Figure 2F).

To determine the distribution pattern of the identified Ψ sites in poly-A RNA, we mapped the RNA reads to the transcriptome reference sequence (gencode.v43.transcripts.fa) and observed that within mRNA, Ψ sites were highly enriched in the coding sequence (CDS) and 3′ untranslated region (3’UTR) (Figure 2G), as reported previously using CeU-Seq^5^, Bid-seq^13^ and Nanopore sequencing methods^11,16^. In addition, we found low levels of Ψ distribution in retained introns (4.46%) and long non-coding RNA (1.14%) (Figure 2G). The limited detection of Ψ sites in the 5′ UTR could be due to a lower level of coverage of the 5′ UTR compared to CDS and 3’UTR as nanopore sequencing moves in the 3’ to 5’ direction. To address this issue, we generated a metatranscript plot using R2D tools^20^ to compare the proportion of significant Ψ sites along mRNA transcripts, which showed comparable Ψ density distribution in the 5’UTR, CDS and 3’UTR (Figure 2H).

Martinej et al.^21^ recently reported that Ψs are enriched in proximal introns (within 500 nucleotides of splice sites). In agreement with the report, we found that the Ψ sites in BE(2)-C poly-A RNA were enriched around 200 nucleotides from the exon-exon junctions (Figure 2I).

### Transcriptome-wide identification and quantification of PUS7-dependent Ψ sites in poly-A RNA

We applied our method to map and quantify PUS7-dependent Ψ sites in BE(2)-C transcriptome. We isolated poly-A RNA from control (shGFP) and PUS7 knockdown (shPUS7-33) BE(2)-C cells (Supplemental Figure 1) and prepared libraries for direct RNA sequencing. Two independent shGFP RNA libraries produced a total of ∼2.254 million reads, with mean read quality of 9.9 and an average N50 read length of 1,124 bases (defined as the shortest read length needed to cover 50% of the sequenced nucleotides). Two independent shPUS7 RNA libraries produced a total of 1.44 million reads, with mean read quality of 10.0 and an average N50 read length of 1,351 bases. The median RNA read lengths were 702 and 789 bases for shGFP and shPUS7-33 samples, respectively (Figure 3A and Supplemental Table 3).

**Figure 3.**
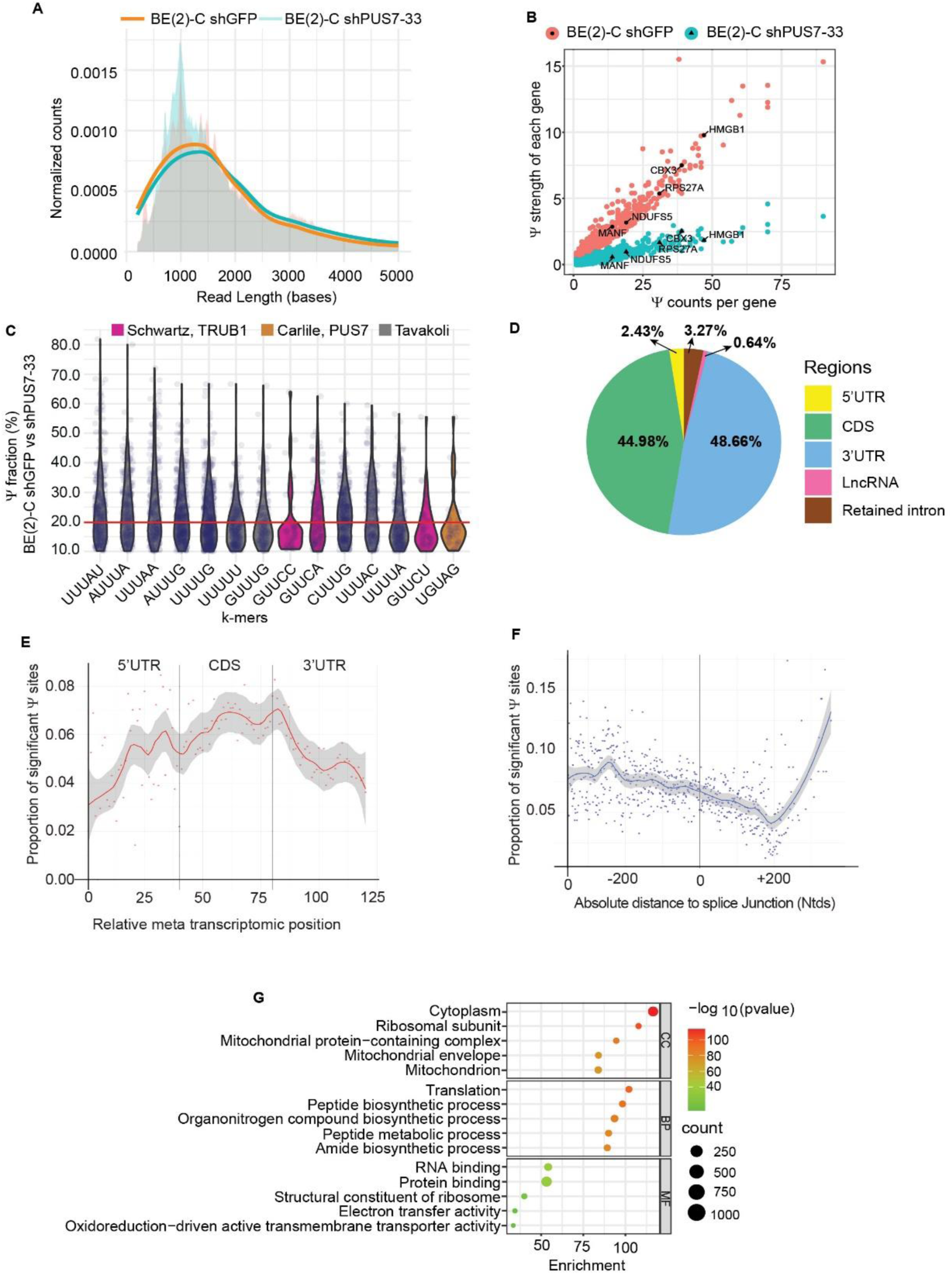
Transcriptome-wide identification and quantification of PUS7-dependent Ψ sites in BE(2)-C cells. **(A)** Normalized counts of read lengths for direct BE(2)-C shGFP (orange) versus BE(2)- C shPUS7-33 (blue) poly-A RNA. (**B)** Scatter plot showing that PUS7 knockdown reduced the Ψ strength of Ψ-modified genes, with labeled representative genes (MANF, NDUFS5, CBX3, RPS27A, and HMGB1). **(C)** K-mer frequency of the most represented and frequently detected PUS7-dependent Ψ sites with high confidence. **(D)** Distribution of PUS7-dependent Ψ sites in mRNA regions, long non-coding RNA, and retained intron. **(E)** MetaTranscript plot showing the proportion of PUS7-dependent Ψ along relative positions in the 5’UTR, CDS, and 3’UTR of mRNA. (**F**) MetaJunction plot showing PUS7-dependent Ψ sites relative to their distances to the closest downstream (left) and upstream (right) exon-exon junction. (**G**) GO analysis of the 1,246 PUS7 target mRNAs. CC, cellular component; BP, biological process; MF, molecular function.

To identify PUS7-dependent Ψ sites, we compared U-to-C basecalling errors at uridine sites in shPUS7 poly-A RNA with Ψ sites in shGFP poly-A RNA that were identified as described above (shGFP BE(2)-C vs. HeLa IVT). Employing the two controls (shGFP BE(2)-C and HeLa IVT), we aimed to enhance the precision of our analysis and eliminate potential false positives in calling PUS7-dependent Ψ sites. We identified 8,624 putative PUS7-dependent Ψ sites within protein-coding mRNA, with Ψ stoichiometries ranging from 10.00% to 88.88% and a median Ψ stoichiometry of 16.78%. Additionally, we identified 294 putative PUS7-dependent Ψ sites within retained introns, with Ψ stoichiometries spanning 10.22% to 68.42% and a median Ψ stoichiometry of 18.08%, and 58 putative PUS7 Ψ sites in long non-coding RNA, exhibiting Ψ stoichiometries between 10.28% and 52% and a median Ψ stoichiometry of 18.18% (Supplemental Table 4). Interestingly, single-read analysis revealed a subset of BE(2)-C mRNAs contain multiple PUS7-dependent Ψ sites within each mRNA molecule. We quantified the overall Ψ modification level in each gene by calculating its Ψ-strength (sum of Ψ fractions at all the Ψ sites within a single gene)^13^. PUS7 knockdown resulted in a global reduction in Ψ-strength of PUS7 target mRNAs (Figure 3B).

Analysis of k-mer frequency and representation for the PUS7-dependent Ψ sites revealed that the previously reported HeLa^11^ Ψ k-mers UUUAU, AUUUA and UUUAA were the most represented, whereas the Ψ k-mers UUUUG, AUUUG and CUUUG were the most frequently detected. The median Ψ stoichiometry of the top 14 k-mers was 19.79% while the median stoichiometry of transcriptome-wide PUS7-dependent Ψ sites was 16.78% (Figure 3C).

As presented above for transcriptome-wide Ψ sites in shGFP BE(2)-C mRNAs (Figure 2G), PUS7-dependent Ψ sites also showed a similar distribution pattern, with ∼48.66% of the Ψ sites in the 3′UTR, 44.98% in the CDS, and 2.43% in the 5′UTR (Figure 3D). Again, the metatranscript plot for PUS7-dependent Ψ sites showed comparable density enrichment in the 5’ UTR, CDS, and 3’UTR (Figure 3E). We also found that PUS7-dependent Ψ sites were enriched around 200 nucleotide proximal from the exon-exon junctions (Figure 3F).

We identified 8,624 putative PUS7-dependent Ψ sites in 1,246 unique protein coding genes. To interrogate the functions of these genes, we performed Gene Ontology (GO) analysis, which revealed the PUS7 target genes are involved in ribosome biogenesis, translation, RNA and protein binding, and mitochondrial energy metabolism (Figure 3G and Supplemental Table 5). Further, we identified 57 Ψ sites in 13 unique long non-coding RNAs. GO analysis found highly significant enrichment of GO terms corresponding to RNA processing (Supplemental Table 6).

### Comparative analysis of rRNA pseudouridylation in HeLa and BE(2)-C cells

Marchand et al. have identified 104 Ψ sites in HeLa rRNA using HydraPsiSeq^22^. To further validate our nanopore sequencing method, we performed nanopore sequencing of rRNA libraries from BE(2)-C cells and obtained 367,424 reads from two BE(2)-C shGFP replicates and 177,515 reads from two BE(2)-C shPUS7-33 replicates. We mapped the U-to-C basecalling error of each uridine base in 5.8S, 18S, and 28S rRNAs and highlighted the previously annotated Ψ sites from HeLa^22^ (Figure 4A-C). We calculated the U-to-C basecalling error for all the 1,510 rRNA uridine bases using direct RNA Nanopore sequencing. Among them, 1,250 bases showed less than 10% U-to-C basecalling errors and were therefore excluded from the analysis (Supplemental Table 8). Out of the remaining 260 uridine bases that showed greater than 10% U-to-C basecalling errors, we identified 97 uridine bases (93.3% of the 104 uridine bases) that were previously annotated as Ψ sites in HeLa rRNA using HydraPsiSeq^22^ (Supplemental Table 9). The remaining 7 bases (6.7%) with less than 10% U-to-C basecalling errors in BE(2)-C could be due to cell-type specificity. This result underscores the high sensitivity of nanopore direct RNA sequencing in detecting previously annotated Ψ sites. Notably, we did not observe any enrichment of specific k-mers associated with these 97 Ψ sites in rRNA (Figure 4D).

**Figure 4.**
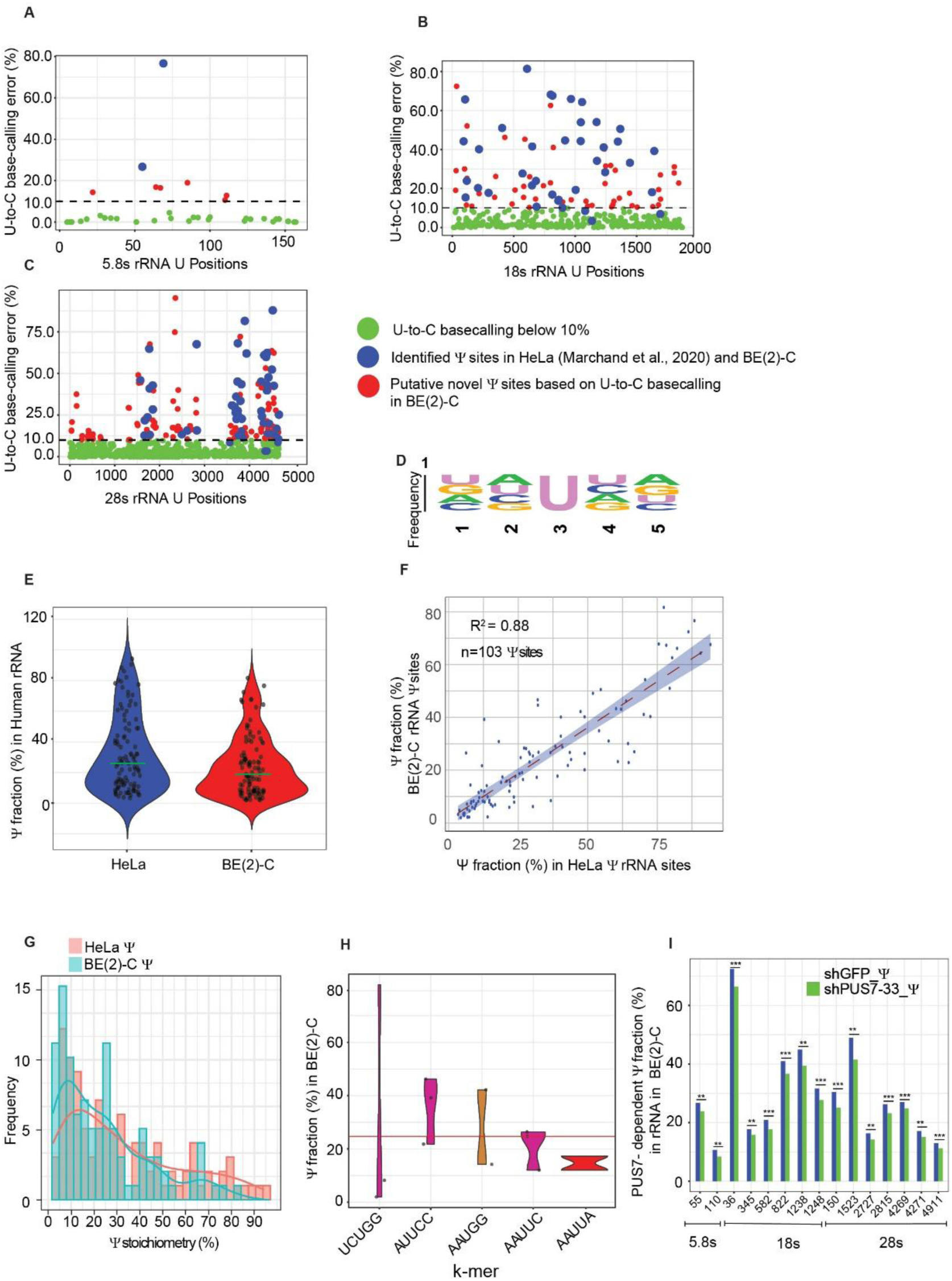
Comparative analysis of rRNA pseudouridylation in HeLa and BE(2)-C cells. **(A-C)** Identification and quantification of Ψ sites in 5.8S, 18S and 28S rRNAs from BE(2)-C and HeLa cells. (**D)** Motif analysis of k-mers for Ψ sites shared between BE(2)- C and HeLa rRNAs. **(E)** Distribution of Ψ stoichiometry in HeLa and BE(2)-C rRNAs identified by nanopore direct RNA sequencing. **(F-G)** Scatter plot (**F**) and bar plot (**G**) showing the distribution of Ψ stoichiometry for the 103 Ψ sites common to both BE(2)-C and HeLa rRNAs. **(H)** Most represented Ψ k-mers identified in BE(2)-C and HeLa cells. (**I**) PUS7 dependent Ψ sites in BE(2)-C rRNAs.

Additionally, we compared our BE(2)-C rRNA pseudouridylation data with the HeLa data that were also generated using nanopore direct RNA sequencing^11^. Among the 103 common Ψ sites identified in the 5.8S and 18S rRNAs of both HeLa and BE(2)- C cells, 65 sites (63.1%) exhibited more than 10% U-to-C base-calling errors in both cell lines, indicating a high degree of conservation for these Ψ sites across cell lines. The median Ψ levels across all Ψ sites in the 5.8S and 18S rRNAs were 25.87% for HeLa and 18.77% for BE(2)-C, with a correlation coefficient of 0.88 (Figure 4E-G). (Supplemental Table 10). Also, we found that the UCUGG k-mer is the most represented k-mer among rRNA Ψ sites identified in both HeLa^11^ and BE(2)-C cells (Figure 4H).

Finally, we identified 15 Ψ sites where the U-to-C basecalling error was significantly reduced after PUS7 knockdown. Notably, 8 of these 15 sites were previously identified as Ψ sites in HeLa rRNA (Figure 4I and Supplemental Table 11).

Collectively, our rRNA Ψ analysis demonstrates that nanopore direct RNA sequencing can reliably identify previously annotated Ψ sites.

### PUS7-dependent pseudouridylation represses m^6^A modification in mRNA

Huang et al. reported recently that transcripts with higher numbers of Ψ sites have fewer m6A sites, while those with more m^6^A sites exhibited reduced levels of Ψ modification, suggesting an antagonistic relationship between Ψ and m6A in mRNA^23^. In agreement with the finding, our dot blot analysis showed a 1.6-fold increase in m^6^A levels following PUS7 knockdown (Figure 5A). To further investigate the impact of PUS7-dependent pseudouridylation on m^6^A modification, we analyzed the same nanopore sequencing data for m^6^A levels in poly-A RNA from BE(2)-C shGFP and shPUS7 cells using two independent pipelines, CHEUI^20^ and m6anet^24^. We divided the PUS7-dependent Ψ data into ten fractions, ranging from 0% to 100% in 10% pseudouridylation stoichiometry increments, which revealed a significant reduction in Ψ levels across all fractions following PUS7 knockdown (Figure 5B and Supplemental Table 12). The CHEUI and m6anet tools identified 4,537 and 1,223 m6A sites within m^6^A DRACH motifs, respectively, in BE(2)-C shGFP and shPUS7 mRNA samples. For the m6anet analysis, we used a probability threshold of 0.9 to select m^6^A sites, ensuring high confidence in the identified sites. Both analyses showed a significant increase in the numbers of m^6^A sites in higher m^6^A stoichiometry fractions after PUS7 knockdown (Figure 5C-D and Supplemental Table 13-14), providing further evidence that PUS7 knockdown increases the global m^6^A levels in mRNA.

**Figure 5.**
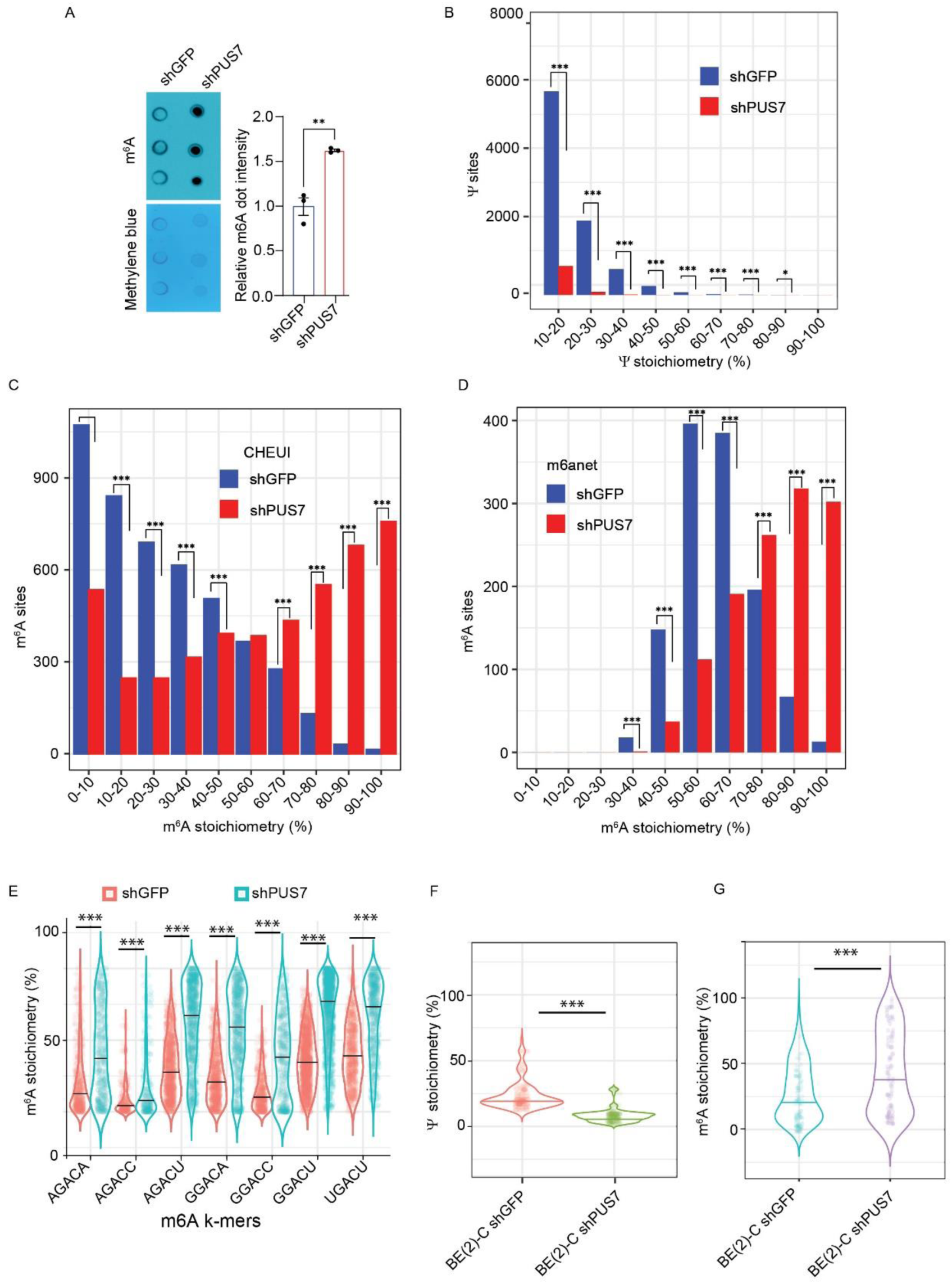
PUS7-dependent pseudouridylation represses m6A modification in mRNA (**A**) m^6^A dot blot of 100 ng poly-A RNA from BE(2)-C shGFP and shPUS7 cells (n = 3 biological replicates). Methylene blue blot serves as loading control. **(B)** Bar graph showing the numbers of Ψ sites across Ψ stoichiometry fractions following PUS7 knockdown. **(C-D**) Bar graphs showing the numbers of m^6^A sites across various m^6^A stoichiometry fractions following PUS7 knockdown analyzed by CHEUI (**C**) or m6anet (**D**) pipelines. **(E)** Violin plot showing m^6^A stoichiometry fractions for various m^6^A DRACH k-mers following PUS7 knockdown identified by CHEUI. (**F-G**) Violin plots showing the effects of PUS7 knockdown on Ψ (**F**) and m^6^A (**G**) stoichiometry when they co-occur on the same transcripts. *p < 0.05, ***p <0.001.

Next, we examined if the antagonistic relationship between m^6^A and Ψ modifications in mRNAs is specific to certain DRACH motifs. We compared m^6^A stoichiometry between BE(2)-C shGFP and shPUS7 across the seven most common DRACH m^6^A motifs and observed an average increase of 2.60 ± 0.74-fold in m^6^A stoichiometry levels following PUS7 knockdown (Figure 5E).

We further investigated whether this antagonistic relationship between m^6^A and Ψ persists when they co-occur in the same transcripts. We identified a total of 104 m^6^A and Ψ sites on 45 unique mRNA transcripts containing any of the three previously verified Ψ k-mers (UGUAG, GUUCA, and GUUCC) and m^6^A DRACH motifs and then analyzed m^6^A levels using CHEUI. Consistently, we found that as Ψ levels decreased, m^6^A levels increased (Figure 5F-G and Supplemental Table 15).

Collectively, the data presented above suggest that PUS7-dependent pseudouridylation has a significant role in regulation of m^6^A modification in mRNA.

### PUS7-dependent pseudouridylation represses m^5^C modification in mRNA

Considering above findings on m^6^A, we investigated the relationship between m^5^C and Ψ modifications in mRNA. Dot blot analysis revealed a 2.3-fold increase in m^5^C levels following PUS7 knockdown (Figure 6A). Using CHEUI, we identified 37,276 m^5^C sites in BE(2)-C shGFP and shPUS7 poly-A RNA (Supplemental Table 16). Our analysis was focused on the 3,014 m^5^C sites linked to the most frequently occurring k-mers: GACUG (2.3%), GACUU (2.2%), GACUC (1.8%), GACAC (1.0%), and GACAU (0.9%) (Supplemental Table 17). We measured the levels of m^5^C in 10% gradient fractions of control and PUS7 knockdown samples, which revealed that PUS7 knockdown increased the numbers of m^5^C sites in poly-A RNAs with higher m^5^C stoichiometries (Figure 6B and Supplemental Table 17). Also, we found that PUS7 knockdown resulted in a significant increase in m^5^C stoichiometry levels in these k-mers (Figure 6C).

**Figure 6.**
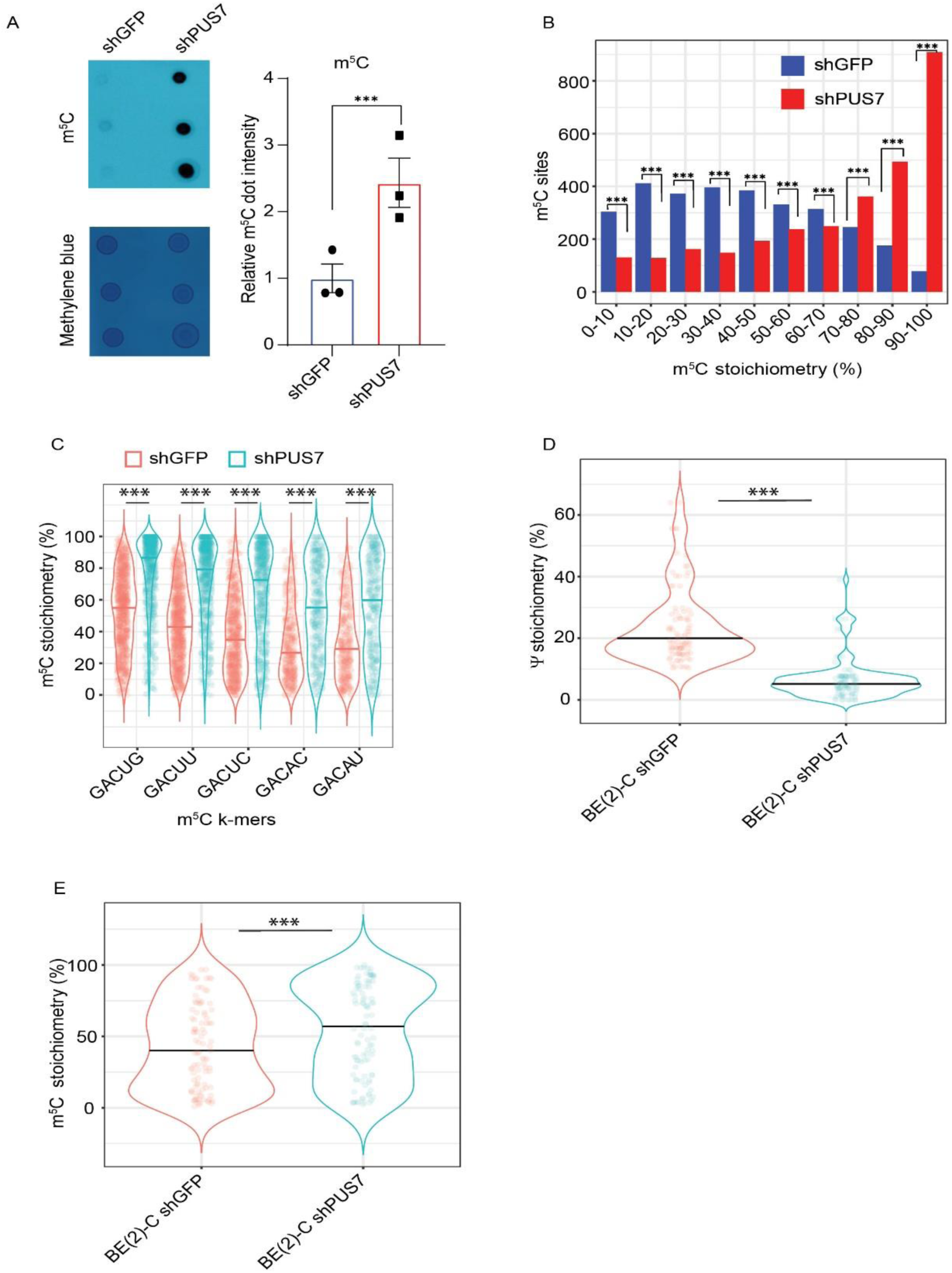
PUS7-dependent pseudouridylation represses m^5^C modification in mRNA **(A)** m^5^C dot blot of 100 ng poly-A RNA from BE(2)-C shGFP and shPUS7 cells (n = 3 biological replicates). Methylene blue blot serves as loading control. (**B**) Bar graph showing the numbers of m^5^C sites across various m^5^C stoichiometry fractions following PUS7 knockdown analyzed by CHEUI. (**C**) Violin plot showing PUS7 knockdown on m^5^C stoichiometry for the most common m^5^C k-mers analyzed by CHEUI. **(D-E)** Violin plots showing the effects of PUS7 knockdown on Ψ (**D**) and m^5^C (**E**) stoichiometry when they co-occur on the same transcripts. ***p <0.001.

We further investigated if this antagonistic relationship between Ψ and m^5^C persists when they co-occur on the same transcripts. To maximize accuracy, we selected transcripts containing experimentally verified Ψ k-mers (UGUAG, GUUCA, and GUUCC) and the 5 most frequent m^5^C k-mers (GACUG, GACUU, GACUC, GACAC, and GACAU). We identified a total of 97 m^5^C and Ψ sites on 53 unique mRNA transcripts that meet this criterion. PUS7 knockdown resulted in a 3.7-fold decrease in Ψ levels in these transcripts (Figure 6D), which was accompanied by a significant increase in m^5^C levels, with the median stoichiometry increasing from 38.5% in shGFP cells to 55.6% in PUS7 knockdown cells (Figure 6E, Supplemental Table 18).

Taken together, these data suggest that PUS7-dependent pseudouridylation has a significant role in regulation of m^5^C modification in mRNA.

## Discussion

We have developed NanoPsiPy for processing nanopore direct RNA long-read sequencing data to identify Ψ at individual RNA molecule and whole transcriptome levels with location information and stoichiometry quantification. NanoPsiPy abstracts U-to-C basecalling errors to detect and quantify transcriptome-wide Ψ modifications between any two samples using comparison paradigm to minimize false positives. By treating in vitro RNA oligos with wild-type and mutant (D294A) PUS7, our study demonstrated that U-to-C basecalling is the sole characteristic enabling the identification of Ψ in nanopore direct RNA sequencing data. NanoPsiPy weigh the pooled sample analysis to identify transcriptome-wide Ψ, comparing the average of U- to-C errors between two conditions to minimize false positives due to low coverage that is an issue with direct mRNA sequencing.

Quantitative MS analysis estimated that the Ψ/U ratio is approximately 0.2-0.6% in human mRNA^5^, which is similar to the ratio of m^6^A/A at 0.15-0.6% found in the human transcriptome^25–27^. Using NanoPsiPy, we identified 39,756 Ψ sites in poly-A RNA from the human neuroblastoma cell line BE(2)-C, which again is comparable to the 42,116 m^6^A sites detected in poly-A RNA from the human mammary epithelial cell line HMEC using MINES analysis of nanopore direct RNA sequencing data^28^. Recently, Tavakoli et al. reported the identification of 1,691 Ψ sites in HeLa poly-A RNA using PsiNanopore that also relies on U-to-C basecalling errors in direct RNA sequencing ^11^. Our analysis identified 517 Ψ sites that are shared between BE(2)-C and HeLa cells, suggesting that a common set of poly-A RNAs are targeted by pseudouridylation in human cancer cell lines of different tissue origins. On the other hand, we identified significantly more Ψ sites in BE(2)-C poly-A RNAs. We suspect that a major reason for the differences in Ψ locations and numbers between HeLa and BE(2)-C cells is the cell-type specific transcriptome profiles because modifications in highly expressed RNAs are most likely to be detected. BE(2)-C cells are of neuronal origin with amplification of the MYCN oncogene, a member of the MYC family of transcription factors that regulates RNA expression of thousands of genes ^29,30^. Thus, NanoPsiPy is highly effective in comprehensively identifying Ψ sites and quantifying their levels in the whole transcriptome.

It was reported recently that the median Ψ stoichiometry in HEK293T mRNA is approximately 10%, with 80% of the Ψ sites showing Ψ levels < 20% and 10% showing > 40% ^14^. In agreement, our NanoPsiPy analysis revealed ∼66% of the Ψ sites in BE(2)- C poly-A RNA having modification levels at 10-20%, 23% at 20-30%, and 11% at > 30%. Interestingly, the median methylation level of m^5^C sites in mRNA is in the range of 15-18%. Across various tissues or cell types, the majority (62–70%) of m^5^C sites exhibited low methylation levels (<20%), and 8-10% showed moderate to high methylation levels (>40%)^31^. Collectively, these findings support the notion that RNA modifications occur at relatively low stoichiometric levels across the entire transcriptome.

Previous studies suggest that Ψ distribution in mRNAs is overrepresented in CDS and 3′UTR and underrepresented in 5’UTR ^11,13^. However, our analysis indicates that Ψ is proportionally present in all regions, showing no significant bias. In principle, this broad distribution endows Ψ modifications with the ability to influence many aspects of mRNA metabolism and function. In addition, we identified 1,246 PUS7 target mRNAs in the *MYCN*-amplified neuroblastoma cell line BE(2)-C and GO analysis suggest that they are involved primarily in ribosome biogenesis, translation, and mitochondrial energy metabolism. These findings lay the foundation for further investigation of the functional significance of PUS7-dependent pseudouridylation in neuroblastoma and other MYC-driven cancers.

A major finding of our study is that PUS7 knockdown markedly reduced Ψ levels while increased m^6^A and m^5^C levels in the same transcripts, indicating a crucial role of PUS7-dependent pseudouridylation in regulation of other RNA modifications. The underlying molecular mechanism remains to be elucidated. Our findings support a recent report of an antagonistic relationship between Ψ and m^6^A ^23^ and provide direct evidence that a decrease in Ψ levels can cause an increase in m^6^A and m^5^C levels.

In summary, we have developed a method for quantitative profiling of transcriptome-wide Ψ sites, enabling the identification and quantification of PUS7- dependent Ψ sites at single mRNA molecule resolution with functional implications. We anticipate that NanoPsiPy will facilitate the identification and functional characterization of pseudouridylation and aid in studying cross-talk between RNA modifications.

## Material and methods

### Cell culture and PUS7 knockdown

Neuroblastoma BE(2)-C) cells were cultured in DME/F-12 1:1 (HyClone SH30023) supplemented with 10% Fetal Bovine Serum (Fisher Scientific FB12999102) and 1% Penicillin-Streptomycin (Lonza 17602E). The PUS7 knockdown lentiviral construct shPUS7-33 (TRCN0000061933) was obtained from Sigma-Aldrich. BE(2)-C cells were infected with lentiviruses expressing shGFP or shPUS7-33, and PUS7 knockdown was confirmed by immunoblotting using rabbit anti-PUS7 (1:1000, Thermo Fisher PA5- 54983).

### Poly-A RNA isolation, library preparation, and nanopore sequencing

Total RNA was obtained through TRIzol extraction, and poly(A) mRNA was isolated using the NEBNext Poly(A) mRNA Magnetic Isolation kit (NEB E7490L). The RNA library for direct RNA sequencing was prepared from ∼500 ng of poly-A RNA according to the Oxford Nanopore technology (ONT) direct RNA sequencing protocol version direct-rna-sequencing-sqk-rna002-DRS_9080_v2_revS_14Aug2019-minion. For sequencing, the ONT FLO-MIN106D (R9.4.1) flow cell was primed according to the manufacturer’s protocol, and the elute was mixed with an RNA running buffer (RRB) and loaded onto the flow cell. Sequencing was performed using MinKnow (version 19.12.5). Each flow cell was sequenced for 24-72 hours per each run, depending on flowcell exhaustion rate. Two replicates from different passages and different flow cells were used for each biological replicate. To study rRNA pseudouridine modifications, we prepared a direct RNA sequencing library from total RNA, similar to the approach used for poly-A mRNA sequencing. We first added a poly-A tail to the total RNA and then sequenced it using Flongle flow cells.

### PUS7 immunoprecipitation and purification

All immunoprecipitation steps were performed at 4°C following the protocol of Kundu et al. with some modifications^32^. Briefly, 293FT cells expressing either wild-type (WT) or mutant (D294A) PUS7 (myc-tagged) were lysed in the buffer containing 50 mM HEPES pH 7.5, 130 mM NaCl, 0.3% NP-40, 1 mM EDTA, and 1× protease/phosphatase inhibitor (Thermo Fisher 78440). Approximately 3 mg of protein lysates were mixed with protein A/G-magnetic beads (Thermo Fisher 88802) and 2 µg of either anti-Myc or control IgG antibody and rotated for 16 hours at 4°C. Subsequently, the magnetic beads were pelleted and washed for three times in the buffer containing 20 mM Tris-HCl at pH 8.0, 130 mM NaCl, 0.3% NP-40, and 1 mM EDTA. Then myc-PUS7 (WT or D294A) was eluted from the beads using myc-peptide (Protein Tech 2yp) following manufacture’s protocol. Finally, using 30 KDa ultracentrifugal filter, the eluted proteins were concentrated, and buffer exchanged in the 1X pseudouridylation buffer (100 mM Tris-HCL pH-8.0, 100 mM ammonium acerate, 5 mM MgCl2, 0.3 mM EDTA, and 130 mM NaCl). Concentrated proteins were either used immediately or stored in -80°C until use.

### In vitro RNA pseudouridylation^18^

The synthetic 192-nucleotide (nt) MALAT1 DNA (Supplemental Table 7) contains 86 nt upstream and 66 nt downstream of the pseudouridine site (U5614, NR_002819.4). A 22-nt sequence containing the T7 promoter and a 25-nt adapter sequence were added to the 5’ and 3’ ends, respectively. The oligo was obtained from IDT DNA and amplified by PCR using Phusion Polymerase (NEB F630) and purified for in vitro transcription using the MEGAshortscript T7 kit (Invitrogen AM1333). The RNA product was purified using a urea gel, denatured, and refolded. In vitro pseudouridylation was carried out by incubating 15-30 pmol of folded RNA with 1 µg of immunopurified PUS7 or PUS7 D294A. Poly-A tail was added using a Poly (A) tailing kit (Invitrogen AM1350). Library was prepared and loaded into the FLO-MIN106D flow cell for sequencing using MinKnow (version 19.12.5) as described above. Each flow cell was sequenced for 24 h.

### m^6^A and m^5^C analyses

CHEUI^20^ and m6anet^24^ were employed to analyze the same nanopore direct RNA sequencing data from BE(2)-C shGFP and shPUS7 cells to identify and quantify m^6^A levels in transcripts containing DRACH motifs. Likewise, CHEUI was used to analyze the same nanopore data to identify and quantify m5C levels in transcripts. The FASTQ files were aligned to the Gencode transcript reference (gencode.v43.transcripts.fa).

Similar to the Ψ analysis, two replicates from different passages and different flow cells were used for each biological replicate.

### Dot blot analysis

Dot blot was performed with either poly-A RNA or total RNA. The poly-A RNA was isolated from total RNA using a NEB poly-A RNA isolation kit. RNA concentration was measured using a Qubit (Thermo Fisher Scientific). RNA was dotted on a nitrocellulose membrane (Bio-Rad). Once the membrane was dried, RNA was cross-linked to the membrane using 120 mJ/cm2 density of UV light (UVP Hybrilinker HL-2000). The membrane was washed once with 2× SSC solution (Thermo Fisher Scientific), incubated with 1× Blot Stain Blue (MilliporeSigma) for 1-2 minutes, and then rinsed several times with 1× TBST (Tris-buffered saline, pH 7.4, plus 0.1% Tween-20) to minimize the background. After imaging of the blue RNA dots, the membrane was blocked with 5% milk for 1 hour, followed by overnight incubation at 4°C with anti-m^6^A antibody (Cell Signaling; mAb #56593; 1:1,000 dilution) or anti-m^5^C antibody (Proteintech; Cat No. 68301-1; 1:000 dilution). After washing, the membrane was incubated with HRP-conjugated secondary antibody for 1 hour (1:3,000 dilution) and the image was captured using a ChemiDoc Imaging System (Bio-Rad).

### Data analysis

All raw fast5 files generated from nanopore sequencing were uploaded to UAB Cheaha cluster for further analysis. Multi-fast5 files were basecalled in real-time by Guppy (6.4.2) using the high accuracy model with minimum quality score of 7. The fastq files were analyzed using the NanoPsiPy where reads were aligned to the reference file (GRCh38.p14 or gencode.v43.transcripts.fa.) using minimap2 (2.17). The mapped reads were piled up to the human reference sequence by samtools (v1.12). The U-to-C basecalling “error” data were extracted and compared between two conditions using NanoPsiPy (http://github.com/vetmohit89/NanoPsiPy.git). NanoPsiPy tool was built around Nanopore_psu tool available at (https://github.com/sihaohuanguc/Nanopore_psU/)^16^. Finally, data was prepared by applying condition when the ratio of C reads from the total of U+C reads of the control group is greater than the ratio of C reads from the total of U+C reads for the treatment group. After filtering the input data, the Chi-square test is applied to a contingency table constructed from the read counts in different categories (control_C, control_U, treatment_C, treatment_U). Yates’ continuity correction is applied to improve the accuracy of the results when the sample size is limited and p < 0.01 is defined as significant. Gene ontology (GO) analysis was performed using gprofiler2 (https://biit.cs.ut.ee/gprofiler/gost).

### Accession Numbers

All Fast5 files generated in this study have been publicly shared and are accessible through the NIH NCBI SRA under the Bio-Project accession PRJNA961708 (SRR29662300, SRR29662301, SRR29662302 and SRR29662303). The data for two HeLa IVT RNA samples were retrieved from NIH NCBI SRA with the accession numbers SRR23932675 and SRR23929127.

## Supplemental Tables description

**Supplemental Table 1:** Significant Ψ sites identified in BE(2)-C poly-A RNA.

**Supplemental Table 2:** Common Ψ sites identified in BE(2)-C and HeLa poly-A RNA.

**Supplemental Table 3:** Direct RNA Library description of BE(2)-C shGFP and shPUS7- 33.

**Supplemental Table 4:** Significant PUS7-dependent Ψ sites in protein coding, retained RNA and long non-coding RNA of BE(2)-C cells.

**Supplemental Table 5:** Gene ontology analysis of protein coding mRNAs containing PUS-dependent Ψ sites

**Supplemental Table 6:** Gene ontology analysis of long non-coding RNAs containing PUS7-dependent Ψ sites.

**Supplemental Table 7:** RNA oligo sequences.

**Supplemental Table 8:** U-to-C basecalling errors as a proxy for pseudouridine from BE(2)-C rRNA.

**Supplemental Table 9:** rRNA Ψ modifications (BE(2)-C vs. HeLa, Marchand et al., 2020).

**Supplemental Table 10:** rRNA Ψ modifications (BE(2)-C vs. HeLa, Tavakoli et al., 2023).

**Supplemental Table 11:** PUS7-dependent Ψ sites in BE(2)-C rRNA.

**Supplemental Table 12:** Distribution of Ψ Stoichiometry in poly-A RNA from BE(2)-C shGFP and shPUS7-33 cells.

**Supplemental Table 13:** PUS7 knockdown on m^6^A stoichiometry analyzed by CHEUI.

**Supplemental Table 14:** PUS7 knockdown on m^6^A stoichiometry analyzed by m6anet.

**Supplemental Table 15:** Co-occurrence of PUS7 dependent Ψ and m^6^A stoichiometry in the same transcripts from BE(2)-C cells.

**Supplemental Table 16:** m^5^C sites in poly-A RNA from BE(2)-C shGFP and shPUS7 cells identified by CHEUI.

**Supplemental Table 17:** PUS7 knockdown on m^5^C stoichiometry within the most frequent m^5^C k-mers.

**Supplemental Table 18:** Co-occurrence of PUS7-dependent Ψ and m^5^C stoichiometry in the same transcripts from BE(2)-C Cells.

## Acknowledgment section

AK was partly supported by a Concept Award (Award HT9425-23-1-0339) and an Early Career Award (HT9425-23-1-0783) from the Department of Defense.

## Author contributions

M.B. and H-F.D. conceived the study and designed the experiments with contributions from A.K., A.Gibson, A.Gupta, J.D., and S.S. M.B. performed nanopore RNA sequencing and data analysis with assistance from A.Gibson and S.V.R. A.K. performed PUS7 purification. A.Gupta. and J.D. performed cell culture and immunoblotting. M.B. and H-F.D. wrote the paper with contributions from S.S. and J.D. H-F.D. supervised and provided funding for the project. All authors read the manuscript and approved its contents.

---

**Supplemental Figure 1.**
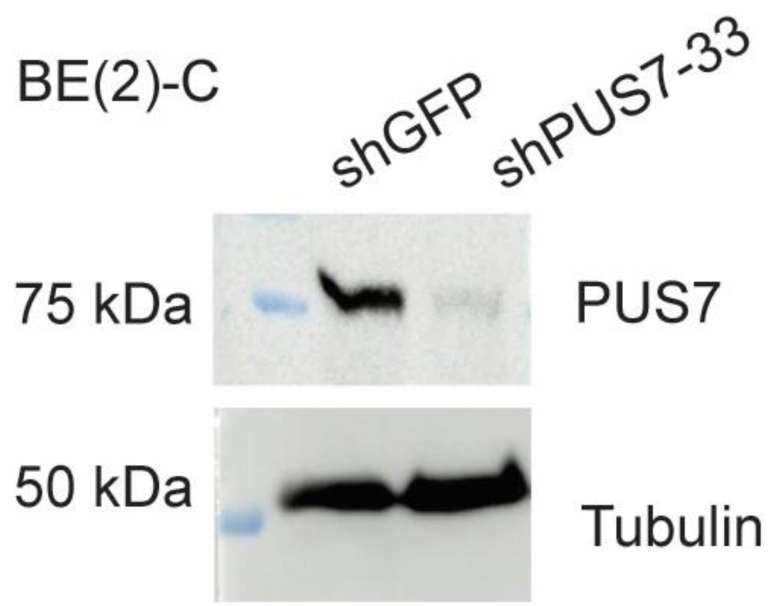
Immunoblotting of PUS7 in BE(2)-C cells expressing shGFP or shPUS7-33.

